# Group and Individual Level Variations between Symmetric and Asymmetric DLPFC Montages for tDCS over Large Scale Brain Network Nodes

**DOI:** 10.1101/2020.06.10.141853

**Authors:** Ghazaleh Soleimani, Mehrdad Saviz, Marom Bikson, Farzad Towhidkhah, Rayus Kuplicki, Martin P. Paulus, Hamed Ekhtiari

## Abstract

Two challenges to optimizing transcranial direct current stimulation (tDCS) are selecting between, often similar, electrode montages and accounting for inter-individual differences in response. These two factors are related by how tDCS montage determines current flow through the brain considered across or within individuals. MRI-based computational head models (CHMs) predict how brain anatomy determine electric field (EF) patterns for a given tDCS montage. Because conventional tDCS produces diffuse brain current flow, stimulation outcomes may be understood as modulation of global networks. Therefore, we developed network-led, rather than region-led, approach. We specifically considered two common frontal tDCS montages that nominally target the dorsolateral prefrontal cortex; asymmetric unilateral (anode/cathode: F4/Fp1) and symmetric bilateral (F4/F3) electrode montages. CHMs of 66 participants were constructed. We showed that cathode location significantly affects EFs in the limbic network. Furthermore, using a finer parcellation of large-scale networks, we found significant differences in some of main nodes within a network, even if there is no difference at the network level. This study generally demonstrates a methodology for considering the components of large-scale networks in CHMs instead of targeting a single region and specifically provides insight into how symmetric vs asymmetric frontal tDCS may differentially modulate networks across a population.

## 1. Introduction

Transcranial direct current stimulation (tDCS) is a non-invasive brain stimulation (NIBS) method based on passing a low-intensity current (e.g. 1-2 mA) from an anode to a cathode electrode at specific scalp locations [1-3]. Neuromodulation effects of tDCS are electrode position (montage) specific and are understood to reflect a combination of different mechanisms of action from the cellular [4-7] to the large-scale functional network scale [8-11].

Despite the effectiveness of tDCS, one major limitation is inter-individual differences in response [12-14]. The intensity and distribution current flow through the brain, reflected in the electric field (EF) magnitude across the brain, is one main reason hypothesized for the individual variations [15-19]. Computational head models (CHMs) are anatomically accurate models, validated by intracranial recording [20, 21], physiological [17], and imaging studies [22, 23] that can estimate EFs as a function of stimulation dose and brain structure [24].

An overarching consideration in optimizing tDCS is dose selection, specifically the electrode montage, current intensity, and duration. Stimulation intensity [25], stimulation duration [26], and electrode characteristics [27] can categorically impart tDCS outcomes. Notably, it has been reported that changing the position of the so-called “return” or “reference” electrode, will impact brain-wide current flow and outcomes of tDCS [2, 28, 29].

The majority of previous computational modeling research to study the effects of electrode location was based on whole-brain or anatomical regions of interest (ROIs) analysis [30-33]. However, brain regions do not operate, or respond to brain stimulation, in isolation, and many distributed regions interact with each other through the brain networks [9, 34-36]. As such, to effectively determine the mechanistic effects of tDCS, rather than concentrating on any given brain regions as currently pursued in modeling studies, the network-led approach for EF comparison in the brain may represent a more productive strategy when comparing stimulation montages.

In a recent study, Fischer et al. suggested that the results of stimulating a brain target may be strengthened or weekend by other brain regions that work together with the targeted region as a network. They hypothesized that modulation of one brain region may impact and be impacted by other regions through the networks and designed a novel multifocal electrode array at the subject-level that stimulate left M1 with excitatory effects, but simultaneously inhibit activity in other remaining nodes of the motor cortex network. Their network approach for placement of the electrodes showed a greater impact on M1 excitability than stimulating just left M1 alone [37]. In another study, through a group-level analysis of computational models, Gomez and his colleagues used a functional atlas for parcellation of the cerebellum. They investigated how the current spreads in the cerebellum networks and reported significant differences between various electrode arrangements for stimulating the cerebellum [38]. However, none of the studies have taken into account the anatomical differences among subjects’ brain variability yet and it remains unclear how much the electrode location can change EF distribution patterns across nodes of large-scale functional networks in a group of subjects with different anatomy.

To bridge this gap, in the present study, we advance a systematic pipeline to investigate EFs inside the brain networks in a group of subjects. We have used structural data from a population with methamphetamine use disorders (MUDs) for generating CHMs. Two of the most common electrode montages for modulating the DLPFC were simulated [39]; an asymmetric “unilateral” montage and a symmetric “bilateral” montage. Our approach allows identifying affected nodes in large-scale functional networks in the standard space and will be insightful for developing hypotheses on the network modulatory effects we should expect. The results of this study provide a better mechanistic understanding of the effects of the montages targeting DLPFC and can be used for guiding the selection of electrode arrangements for future researches.

## 2. Methods and materials

### 2.1. Participants and data acquisition

Participants included 66 subjects (all-male, mean age ± standard deviation (SD) = 35.86 ± 8.47 years ranges from 20 to 55) with methamphetamine use disorder (MUD). All subjects were recruited during their early abstinence from the 12&12 residential drug addiction treatment center in Tulsa, Oklahoma in the process of a clinical trial to measure the efficacy of tDCS on methamphetamine craving (ClinicalTrials.gov Identifier: NCT03382379). None of the participants had a history of neurological disorders, head injury, or other abnormalities demonstrated by sMRI. Written informed consent was obtained from all participants before the scans and the study was approved by the Western IRB (WIRB® Protocol #20171742). After receiving each subject’s written consent, structural MRIs were obtained on a GE MRI 750 3T scanner. High resolution structural images were acquired through magnetization-prepared rapid acquisition with gradient-echo (MPRAGE) sequence using the following parameters: TR/TE = 5/2.012 ms, FOV/slice = 24 × 192/0.9 mm, 256×256 matrix producing 0.938 x 0.9 mm voxels and 186 axial slices for T1-weighted images and TR/TE=8108/137.728ms, FOV/slice=240/2mm, 512×512 matrix producing 0.469×0.469×2mm voxels and 80 coronal slices for T2-weighted images. Gyri-precise CHMs were generated from a combination of high-resolution T1- and T2-weighted MR images for 52 subjects. For a subset of participants (N = 14), T2-weighted MRIs were not available and head models were created only based on T1 images. This study was conducted in accordance with the Declaration of Helsinki and all methods were carried out in accordance with relevant guidelines and regulations.

### 2.2. Creation of head models and tDCS simulation

The flow diagram of the modeling and data extraction pipeline is briefly shown in Fig.1. Head models were generated for all of the subjects to visualize how current flows through the brain in each individual. Modeling of the EF distribution patterns was performed using the standard SimNIBS 3.08 pipeline [40]. Automated tissue segmentation was performed in SPM 12 combined with CAT12 toolbox via the SimNIBS headreco function. The head volume was assigned to six major head tissues including white matter (WM), gray matter (GM), cerebrospinal fluid (CSF), skull, scalp, and eyeballs. Segmentation results were visually examined slice-by-slice to ensure correct tissue classifications. Segmented surfaces were used to create tetrahedral volume meshes and about 3 x 10^6^ elements were assigned to each head model.

**Figure 1:**
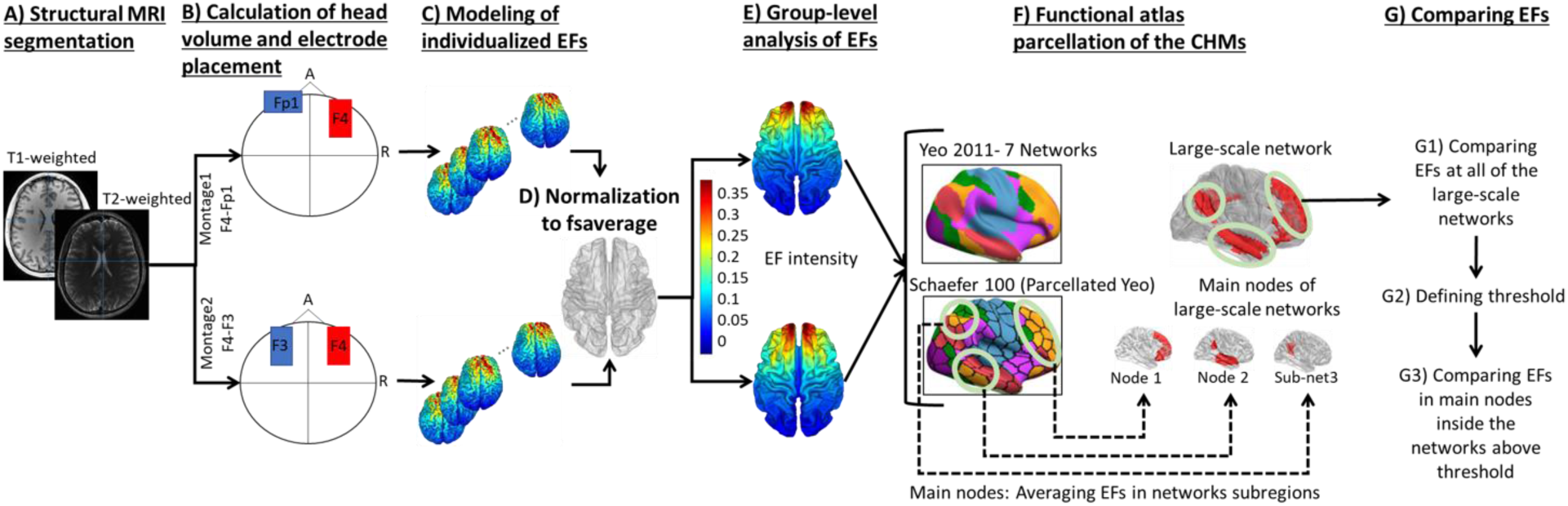
Data extraction flow diagram: (A) Using T1- and T2-weighted MR images for creating CHMs based on segmentation of the structural images into five tissue types. (B) Mesh generation for simulation of two typical conventional electrode montages in stimulation of the DLPFC; anode-cathode in montage1: F4-Fp1 and in montage 2: F4-F3 in 10-20 EEG standard system with 5×7 cm2 electrode pads in 2 mA current intensity (C) Modeling of EF distribution patterns for all of the participants (66 subjects) based on FEM method (D) Normalization of individualized EFs to standard fsaverage space (E) Calculation of strength and normal component of EFs in fsaverage space for group level analyses (F) Using resting-state functional atlas for extracting EFs from brain networks in CHMs (G) Comparing two montages at the level of brain networks by considering a predefined EF threshold inside the networks. Abbreviations: CHMs: computational head models; FEM: finite element modeling; EF: electric field.

Two of the most widely studied tDCS conventional electrode montages for targeting DLPFC were simulated; bipolar asymmetric and bipolar symmetric arrangements [41]. For both montages, 5 cm x 7 cm rectangular pads with 1 mm thickness were generated virtually. In the first montage, as a bipolar asymmetric arrangement (F4-Fp1), the anode electrode was positioned over the F4 electrode location on the standard 10-20 EEG international system with the long axis of the pad pointing towards the vertex of the head. The cathode electrode was positioned over the contralateral eyebrow (Fp1 EEG electrode site also referred to as the supraorbital position) with the long axis of the pad parallel to the horizontal plane. In the second montage, symmetric bipolar (F4-F3), anode and cathode electrodes were respectively placed over F4 and F3 electrode locations on EEG standard system with the long axis of the pads pointing towards the vertex. The locations and directions of the anodes are thus similar in both configurations and the only difference between montages is the position of the cathode electrodes. These arrangements are also referred to as unilateral and bilateral montages in DLPFC stimulation [42, 43] under the assumption the cathode is functionally inert for the asymmetric (unilateral) case.

After electrode placement, the simulations were run with a current amplitude of 2 mA and EFs were calculated based on the finite element method (FEM). To generate CHMs, electrical conductivities were assumed to be constant. The assigned isotropic conductivities were based on the previously reported values; WM = 0.126, GM = 0.275, CSF = 1.654, skull = 0.010, skin = 0.465, eyeballs = 0.5 all in Siemens/meter [44]. The results were visualized using Gmsh [45] and MATLAB (version 2019b, The Math Works, Inc.). The absolute values of the EFs (|EF|) were calculated at each brain node. In order to make the individualized EF maps comparable, the simulation results for all subjects/montages were transformed from the original native space to the standard average surface (‘fsaverage’) of FreeSurfer (http://surfer.nmr.mgh.harvard.edu). In this way, the same coordinates in different subjects correspond to the same location in CHMs. This normalization makes statistics possible at the group-level and allowing for the comparison of the EF distribution patterns between two montages.

### 2.3. Data analysis

Functional atlas parcellation was used for comparing EFs in two montages as distributed over large-scale brain networks. Yeo7-2011 surface-based atlas was used for extracting the topology of the seven resting-state large-scale networks including visual (Vis), somatomotor system (SomMot), dorsal attention (DorsAttn), ventral attention (VentAttn), limbic system (Limbic), executive control network (ECN), and default mode network (DMN) [46]. Since each parcellated network in Yeo7 atlas covered a widely distributed area, we applied Schaefer-100 atlas to CHMs for extracting the finer sub-regions of each functional network [47] to determine the amount of current reaching different parts within a network. We then merged the areas that were placed adjacent to each other to form the main nodes of the networks. Left VentAttn was divided into four main nodes including tempro-occipital-parietal (TempOccPar), frontal-operculum-insula (FrOperIns), prefrontal cortex (PFC; DLPFC node), and medial (Med) nodes. Right VentAttn network parcellated into three sub-networks including TempOccPar, FrOperIns, and Med nodes. The limbic network has two main nodes in each hemisphere including orbito-frontal cortex (Limbic-OFC) and temporal pole (Limbic-TempPole) nodes. The main nodes in the left and right ECN were tempro-parietal (ECN-TempPar), ECN-PFC, and precuneus cingulate (ECN-PcunCing). DMN was also divided into three main nodes in each hemisphere including DMN-TempPar, DMN-PFC, and precuneus posterior cingulate cortex (PcunPCC).

In the next step, the spatial grouped-average EFs as an indicator of modulation intensity were calculated in all of the networks and inter-individual variabilities across the subjects were quantified by standard deviations (SDs). To exclude the networks with low average tDCS-induced EFs from the further analyses in the next steps, we defined a threshold. Maximum EF intensity (an indicator of hotspot inside the brain), 99^th^ percentile, was calculated for each subject and, inspired by [19], 50% of the lowest peak among participants was defined as threshold. Subsequent calculations and comparisons were only performed at networks with mean tDCS EF magnitude higher than this threshold. For statistical analysis, t-tests were used to examine significant differences between the two montages. Averaged EFs in each parcellated regions were compared between the two arrangements. False discovery rate (FDR) correction was used to correct P-values as necessary to overcome multiple comparisons problem and P-value < 0.05 was considered to be significant. All data reported as mean ± standard deviation (SD).

## 3. Results

Surface-based CHMs were simulated for 66 participants with F4-Fp1and F4-F3 montages. Personalized CHMs for F4-Fp1 and F4-F3 montages over all subjects are available in Fig.2 which indicates a visible variation in tDCS induced EFs within a montage among participants.

**Figure 2:**
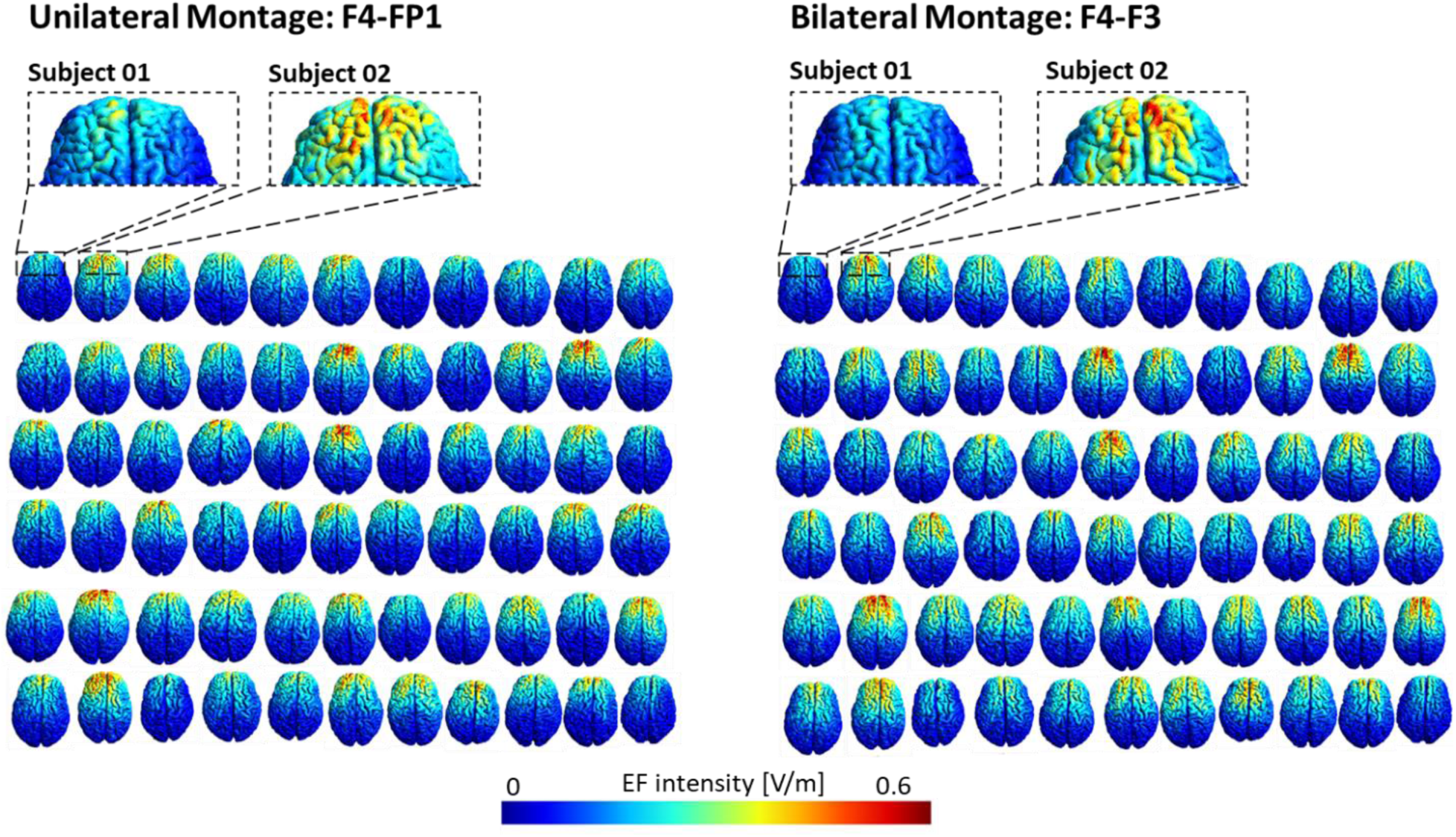
Individualized computational head models: Electric field distribution patterns of 66 participants for DLPFC stimulation with the anode over F4, cathode over the supraorbital area (F4-Fp1 montage) for unilateral montage and anode over F4, cathode over F3 for bilateral montage in subject native space. A current of 2 mA is used in the stimulation.

The spatial global maximum was calculated in the whole brain for each individual as an indicator of the EF hotspot. As shown in Fig.3, a considerable variability for maximum EF is found among subjects but there is no significant difference between F4-Fp1 (0.41 ± 0.07 V/m; range from 0.28 to 0.58) and F4-F3 (0.39 ± 0.07 V/m; range from 0.25 to 0.59) montages in terms of global maximum EF.

**Figure 3:**
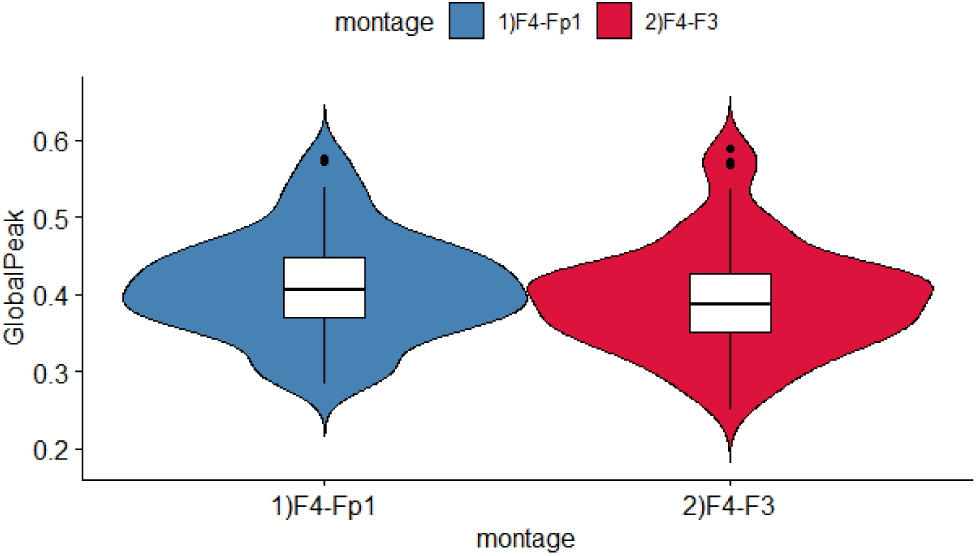
Maximum electric field: Violin plot shows distribution of 99^th^ percentile of the global maximum EF in whole brain analysis for F4-Fp1 (blue) and F4-F3 (red) montages with no significant difference between two montage.

For group-level analysis, the individual EFs were transformed to the standard fsaverage template. Transformation to standard space underestimated the global peak on average by 0.3%, showing that peaks are smoothed a little in transformation to standard space. Group-averaged EF among 66 participants in standard space can be found in Fig.1 part (E) for both montages and it can be found that the highest EF intensity is occurs in the frontal lobe.

### 3.1. Results at the network level

Fig.4 represents the amount of mean EF intensity at seven large-scale brain networks in the right and left hemispheres. The exact amount of mean EF intensity and P values for each network are available in Table.S1 in supplementary materials. The results show that, regardless of montage choice here, the highest EF is produced in limbic system (F4-Fp1: left = 0.1955 ± 0.04, right = 0.155 ± 0.03; F4-F3: left = 0.1540 ± 0.03, right: 0.1544 ± 0.03) and lowest EF can be found in visual network (F4-Fp1: left = 0.0523 ± 0.01, right = 0.0532 ± 0.01; F4-F3: left = 0.0473 ± 0.01, right = 0.0488 ± 0.01) in both montages.

**Figure 4:**
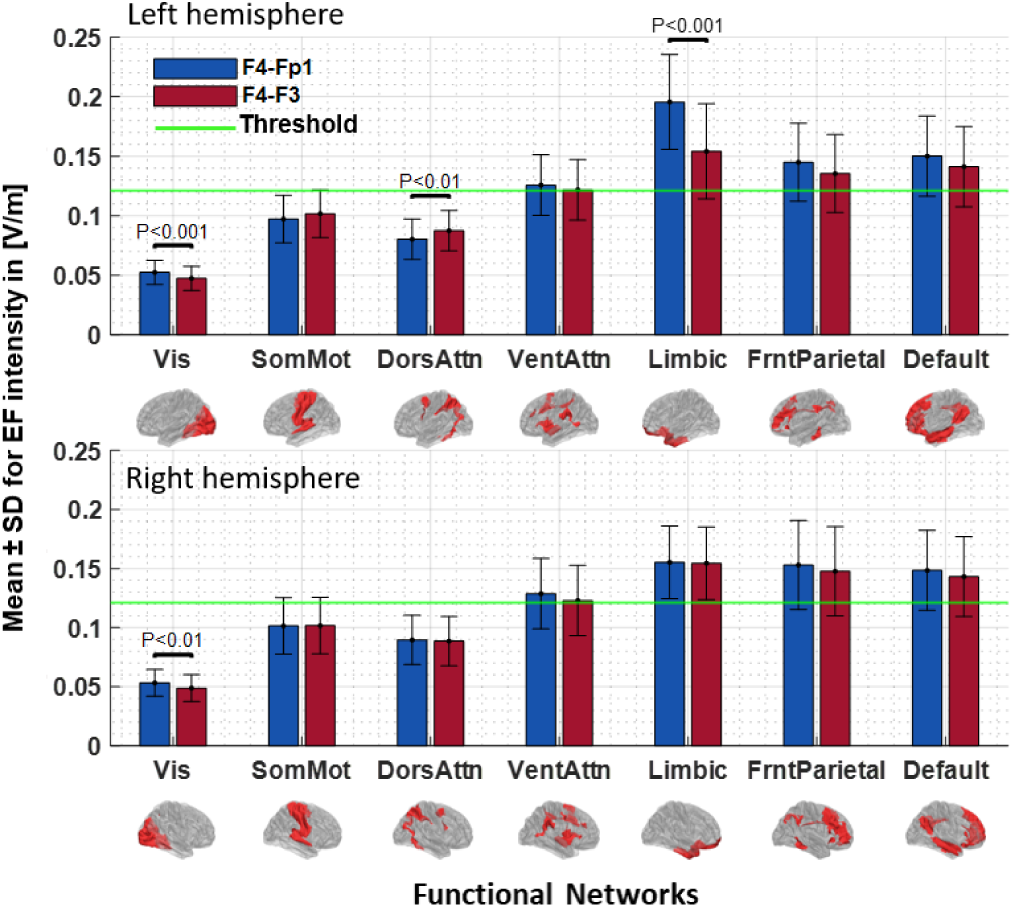
Parcellation of the average EF intensity at a current of 2 mA generate by each montage for the large-scale brain networks. Bars show mean value and error bars show SD of the EF intensity in volt per meter ([V/m]) across the 66 participants in each large-scale network of Yeo7-2011 atlas for F4-Fp1 (blue bars) and F4-F3 (red bars) electrode montages in the left (first row) and right hemisphere (second row). Labels below the horizontal axis determine the name of each network in Yeo7 parcellation and small brains next to the labels represent each network in the fsaverage space. Significant differences between two montages are shown above the bars based on the t-test with FDR correction threshold at P < 0.05. The horizontal green line indicates the EF threshold (50% minimum value of peak EFs in whole-brain analysis across the subjects). Abbreviations: EF: electric field; SD: standard deviation; FDR: false discovery rate; Vis: visual network; SomMot: somatomotor network; DorsAtn: dorsal attention network; VentAtn: ventral attention network; Limbic: limbic system network; ECN: executive control network; DMN: default mode network.

In the left hemisphere, differences between two montages were statistically significant in Vis (P_Corrected_ < 0.01), DorsAttn (P_Corrected_ < 0.01), and Limbic (P_Corrected_ < 0.001) networks. In the right hemisphere, the difference between the two montages was significant only in Vis (P_Corrected_ < 0.01) network. Based on the predefined threshold, as shown with a green horizontal line in Fig.4, tDCS induced EFs in Vis, SomMot, and DorsAttn networks are not above the threshold. Therefore, the left Limbic is the only network that has a significant difference between two montages and EF intensity crosses the threshold in this network. Inspired by [19], we considered 50% of the lowest peak as a threshold (Th = 0.125 V/m) to exclude the networks with low average EFs from further analyses in the next steps. threshold is now represented by the horizontal green line in all of the bar graphs.

The extent of inter-individual variability in the EF was calculated by average induced EF in each network that can be assumed to be normally distributed (Shapiro-Wilk test P > 0.05). Fig.5 shows the corresponding histograms for F4-Fp1 (blue) and F4-F3 (red) montages over 66 participants in the left and right hemispheres. Histograms display inter-individual variability in each subject’s mean EF in the targeted network. We present the distribution of the EF strength in the networks above the threshold and results showed that inter-individual variance is relatively high in all of the networks. Except for the left Limbic, for all other networks, we found relatively similar distribution in both montages.

**Figure 5:**
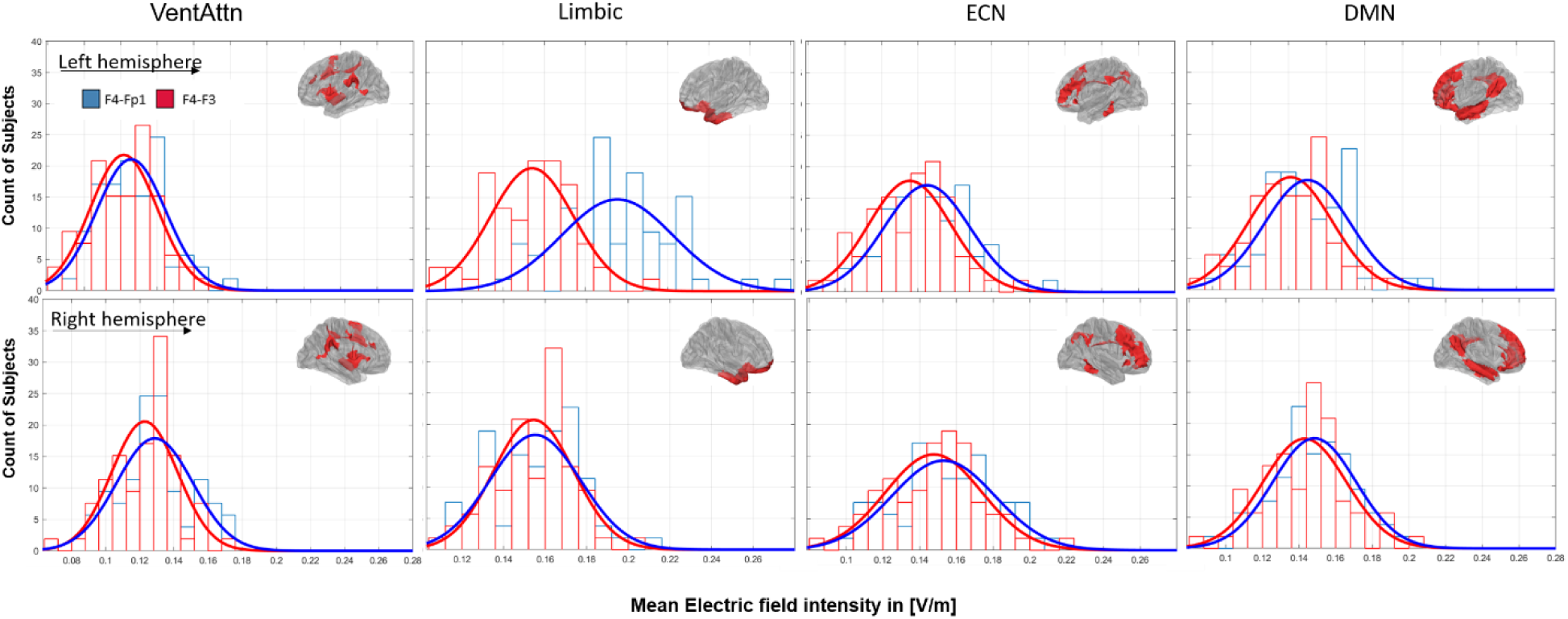
Distribution of the EF in the large-scale brain networks that receive EFs above the threshold. Distribution of the EF intensity in volt per meter ([V/m]) are represented for F4-Fp1 (blue line) and F4-F3 (red line) electrode montages in left (first row) and right (second row) hemisphere. Abbreviations: VentAtn: ventral attention network; Limbic: limbic system network; ECN: executive control network; DMN: default mode network.

Furthermore, inte-individual differences between subjects in the range of EF alterations by changing the electrode montage are represented in Fig.6.

**Figure 6:**
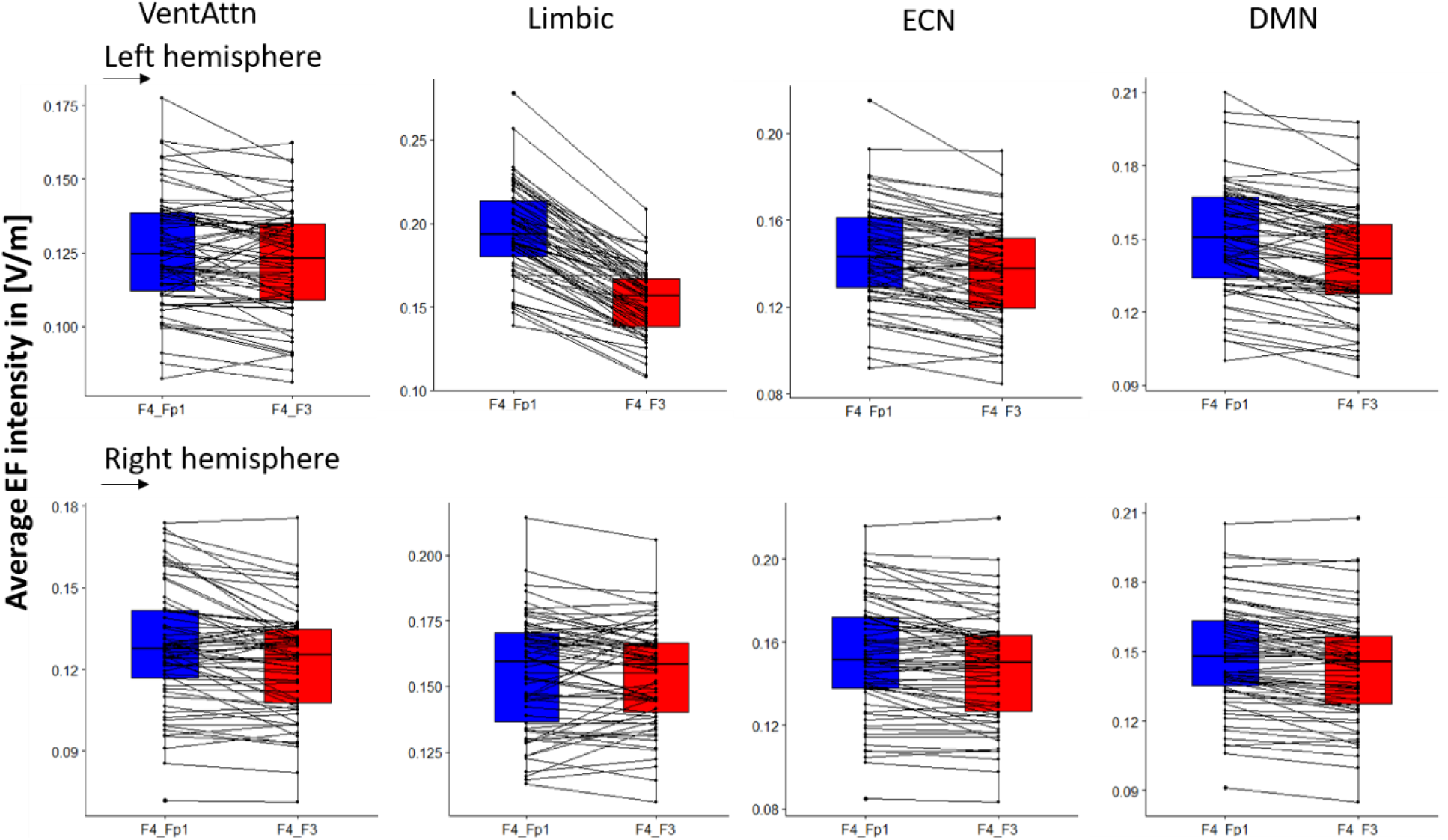
Group-averaged EF intensity for two montages: F4-Fp1 (blue) and F-F3 (red). Average EF intensity calculated for all 66 subjects inside the large-scale brain networks; first row for left and second row for right hemisphere. Results are visualized for the networks with average EF above threshold. Boxplot showing the effects of electrode montage on averaged EF intensity in [V/m] (n = 66 each). Dots represent the data for individual subjects. Abbreviation: VentAtn: ventral attention network; Limbic: limbic system network; ECN: executive control network; DMN: default mode network

### 3.2. Results at the sub-network level

According to the results of the network analysis in the previous step, we showed that applied EFs in the four networks are above the threshold value. In Fig.7, average EFs intensities in the main nodes of these networks are represented separately for each hemisphere. The numerical values of averaged EFs in each node can be found in Table.S2 in the supplementary materials. As indicated in Fig.7, in most areas, EF intensity generated by F4-Fp1 montage is higher than F4-F3. Statistical results indicate that there is no significant difference between the two montages in the nodes with EF above the threshold in the right hemisphere. Nonetheless, in the left hemisphere there are some main nodes with significant differences between two montages including FrOperIns (F4-Fp1: 0.1514 ± 0.03, F4-F3: 0.1385 ± 0.03; P < 0.01) node in VentAttn, OFC (F4-Fp1: 0.2474 ± 0.05, F4-F3: 0.1953 ± 0.04; P < 0.001) and TemPole (F4-Fp1: 0.1342 ± 0.03, F4-F3: 0.1073 ± 0.02; P < 0.001) in Limbic, and ECN-PFC (F4-Fp1: 0.1937 ± 0.04, F4-F3: 0.1661 ± 0.04; P < 0.001) that received EFs above the threshold values.

**Figure 7:**
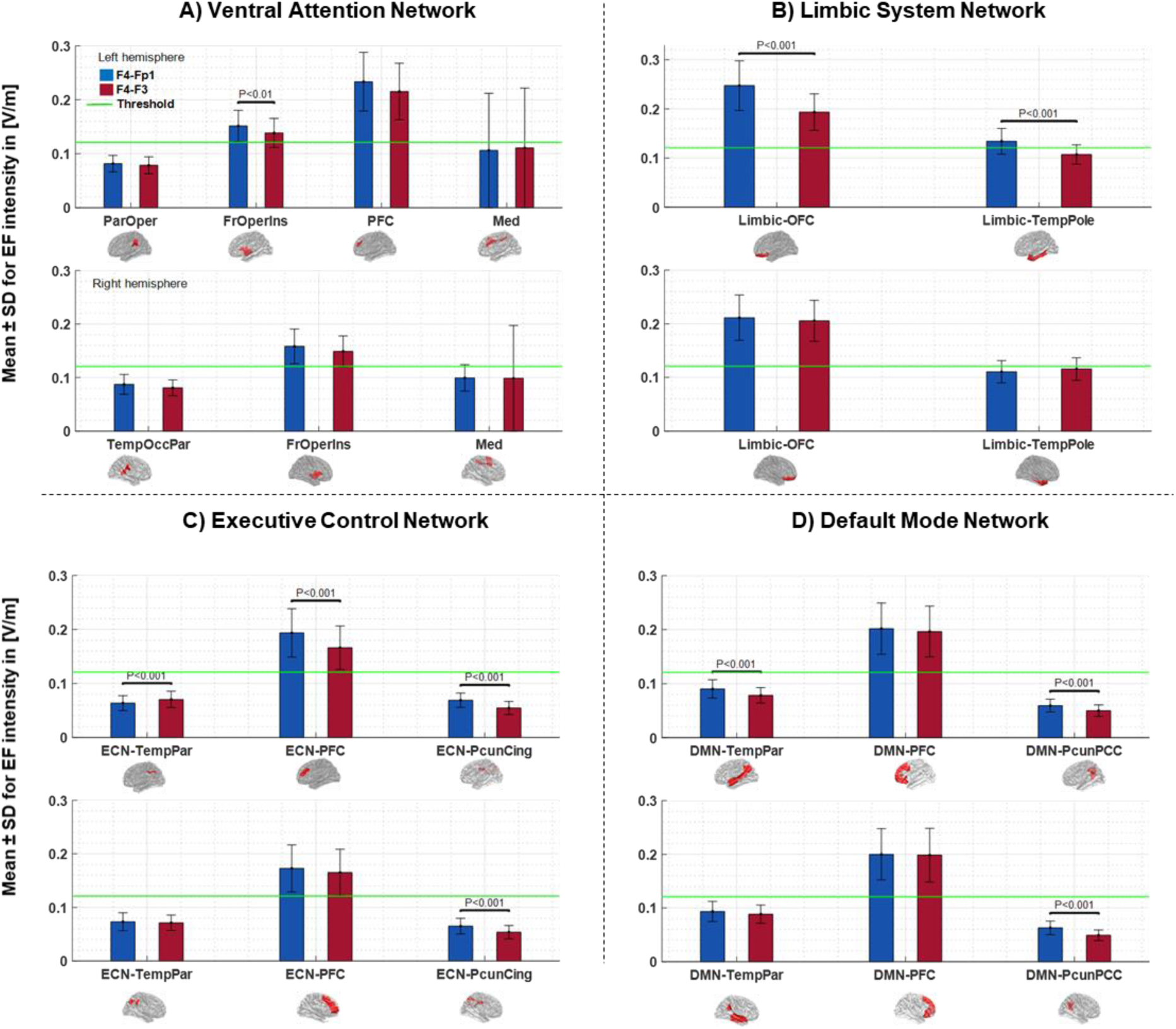
EF intensity in main nodes of the large-scale brain networks at a current of 2 mA generate by each montage. Bars show mean values and error bars represent SDs of the EF intensity in volt per meter ([V/m]) across 66 participants inside the main nodes of large-scale networks for F4-Fp1 (blue bars) and F4-F3 (red bars) electrode montages in the left and right hemisphere of each network. Each network in Yeo7-atlas that received EFs above the threshold including **(A)** Ventral attention network, **(B)** Limbic system network, **(C)** Executive control network, and **(D)** Default mode network parcellated into subregions based on Schaefer2018_100Parcells_7networks atlas and main nodes of each network were extracted by averaging the EFs in the subregions of the main nodes. Labels below the horizontal axis denote the name of main nodes inside the large-scale network and small brains next to the labels represent the spatial location of the main nodes in fsaverage space. Significant differences between two montages are shown above the bars based on t-test with FDR correction threshold at P < 0.05. Horizontal green line indicates EF threshold (50% minimum value of the peak EFs in whole brain analysis across the subjects). Abbreviations: EF: electric field; SD: standard deviation; FDR: false discovery rate. VentAtn: ventral attention network; Limbic: limbic system network; ECN: executive control network; DMN: default mode network. OFC: orbitofrontal cortex; TempPole: temporal pole; TempPar: Tempro-parietal; PFC: prefrontal cortex; PcunCing: Precuneus-cingulate cortex; PcunPCC: Precuneus-posterior cingulate cortex.

## 4. Discussion

In this study, we modeled the network selectivity of two tDCS protocols (F4-Fp1 “unilateral” and F4-F3 “bilateral” montages) that are commonly used for stimulating DLPFC. We tested the individual head models for the electric field (EF) at the group level to examine which parts of the large-scale brain networks are most likely to be stimulated by considering two conventional electrode montages. We showed that placement of the cathode electrode can significantly change EF distribution patterns in specific nodes in large-scale networks even if there is no significant difference in the whole brain network. We found that in the F4-Fp1 montage, the limbic system network in the left hemisphere receives significantly higher electrical dose. Within the network nodes, frontal-operculum-insula nodes in the ventral attention network, orbitofrontal cortex and temporal pole in the limbic system, and DLPFC node in the executive control network significantly receive higher EFs by F4-Fp1 arrangement.

Fig.2 visualizes the inter-individual variability in the EFs. Wide range of variation in the EF distribution patterns over 66 participants supports the previous findings on the importance of considering the individual computational head modeling (CHM) [18, 32, 38, 48]. Individualized head models for a group of subjects with surface-based registration to standard template allowed us to use standard functional atlas parcellation for network-level analysis of the head models.

In this work, we extracted the EF intensity inside the large-scale brain networks and their nodes (Fig.4 and Fig.7). Calculation of EF intensity in brain regions requires a decision about which EF measures (e.g. maximum, average, median) are appropriate to use. Gomez et al. by using network parcellation of the cerebellum reported that in contrast to maximum, functional network analysis becomes important when average EF is used [38].

Our results conform with previous computational modeling studies that cathode location has a significant impact on EF distribution patterns [49, 50]. Previous findings reported that the effects of tDCS depend on the locations of both active and reference electrodes [28, 30].

Our results indicate that among the networks with EF intensity above the defined threshold, the differences between two montages were significant only in the left limbic system. Cathode locations in both arrangements are spatially close but nonetheless cathode over the left orbit is closer to orbitofrontal cortex (OFC) than cathode over left DLPFC (F3). Therefore, F4-Fp1 induced higher EFs than F4-F3 inside this region as the main node of the limbic system. Besides between montage differences, as shown in Fig.4, we also observed within montage variance in terms of averaged EFs inside a network (Fig.5 and Fig.6). For instance, by changing the electrode location from the left supraorbital area to F3 according to the 10-20 EEG electrodes placement standard, there were some subjects whose EFs changed extremely little or oppositely compared to the grouped-average results. For example, changes of EFs in the left limbic system, that showed a significant difference between two montages, range from −2×10^−5^ to 6.9×10^−2^ [V/m] and there is one person (1.5% of total participants) whose EFs changed in opposite direction compared to the grouped-average EFs. However, EF changes in the right limbic system range from −2.8×10^−2^ to 1.7×10^−2^ and there are 19 subjects (28.8% of total participants) whose EFs changed in opposite direction compared to the grouped-average. With respect to previous simulation studies, these inter-individual variations in EF distribution patterns could be explained by different anatomical factors that can change the actual current received by different parts of the cerebral cortex [14, 44, 48]. In fact, current distribution of the tDCS is influenced by microarchitectural features of the brain and variation in cortical anatomy such as gyrus and sulcus patterns can change induced EFs between individuals [51, 52].

EF calculations in the main nodes are depicted in Fig.7 and numerical results can be found in Table.S2 in supplementary materials. According to the brain’s compensatory response to intervention through the neural networks, a simple hypothesis is that modulating multiple nodes of a network would be more effective than modulating a single node in that network [53]. As shown in Fig.7, the intensity of the induced EFs in distinct nodes of a network are different. Grouped-average EF intensity is weak in some nodes in a network since they are located farther from stimulating electrodes. For example, DLPFC, as the main node of executive control network (ECN) was targeted in both montages in our simulations. This node was named ECN-PFC in Fig.7 and received the highest EF inside the network. Statistical results indicated that induced EF by F4-Fp1 in DLPFC was significantly higher than F4-F3. However, EF intensity in other parts of the ECN, precuneus-cingulate and temporal-parietal nodes, is considerably low. This opens potentials for multifocal tDCS montages stimulating all parts of a specific distributed network [37].

We also found that PFC node in DMN network received highest EFs inside the networks and amount of EFs in other nodes of this network is considerably low. The results from an tDCS-fMRI study conducted by Shahbabie et al. demonstrated that application of bilateral DLPFC with the anodal electrode over right DLPFC (same as F4-F3 montage in this study) decreases functional connectivity in DMN especially in posterior parts of the network including middle temporal gyrus, right precuneus, and right PCC [54]. As represented in Fig.7, our calculations showed that temporoparietal and PCC nodes in DMN received a small amount of EFs. Perhaps significant diminishing in functional connectivity of these nodes reported by Shahbbie et al. is related to low EF intensity in these nodes that have not yet been investigated.

Other than ECN-PFC, analysis of the main nodes revealed that, although left limbic system is the only network that shows a significant difference between two montages, however, frontal-operculum-insula node in ventral attention network and orbitofrontal cortex and temporal pole nodes in limbic system all in the left hemisphere are significantly different between these two montages.

These significant differences may become a matter based on stimulation goal. For example, frontal operculum in the ventral attention network is a key node in the cognitive control system. In a combined TMS-fMRI study, it was reported that frontal operculum regulated the level of activity in the occipitotemporal cortex. By applying TMS, they found that stimulation of this region decreased top-down selective attentional modulation in the occipitotemporal cortex [55]. Based on our calculations, F4-Fp1 generated significantly higher EF inside this node of ventral attention network. Therefore, it might be a more effective montage than F4-F3 for stimulating frontal-operculum-insula which is important in manly cognitive tasks including directed attention tasks. However, this is an empirical question for future trials.

As shown in Fig.7, both OFC (within the limbic network) and DLPFC (PFC node in ECN) were stimulated strongly regardless of electrode montage and greater EFs induced by the F4-Fp1 arrangement. These two nodes play important role in different cognitive functions such as decision-making behavior and inhibitory control. A previous tDCS study compared the effects of unilateral (i.e., F4-Fp1 or F3-Fp2) and bilateral (i.e., F4-F3 or F3-F4) DLPFC stimulation on risk taking and decision-making behaviors. Similar to our simulations, they used 5×7 large electrode pads and current strength was 2 mA. They reported that, based on Balloon Analog Risk Task (BART) as a measure of risk-taking propensity, bilateral stimulation of the DLPFC (anode-cathode over F4-F3 or F3-F4), compared to sham, significantly reduced risk-taking behavior. While, no statistically significant difference in risk taking score was discovered between sham and unilateral DLPFC [43]. In a tDCS-fMRI study neural effects of bilateral tDCS (F3-F4) on risk taking behavior during BART task were examined. The authors reported that whole-brain connectivity of right DLPFC (under anodal electrode) correlated negatively with risk taking aversion. It was also reported that OFC activity is negatively correlated with risk and loses in the BART task [56]. However, authors have not investigated the relationships between the induced EFs by unilateral or bilateral montages and neural/behavioral outcomes. Testing the effect of higher levels of EFs in ECN and limbic system generated by unilateral stimulation compared to the bilateral montage, that we found in our simulations, in relationship with neural and behavioral outcomes might give us a better understanding of the underlying mechanism of actions.

Previous research also showed that bilateral stimulation of DLPFC can affect brain in a different way compared to unilateral stimulation during temporal discounting task. Temporal discounting task is a measure of risky decision making, subjective value judgments, and the ability to delay gratification. No change in temporal delay discounting with bilateral tDCS over DLPFC was described by [57]. However, the unilateral DLPFC stimulation (F3-Fp2) modulated temporal discounting rate [58]. Future tDCS studies on cognitive functions critically depend on OFC and DLPFC activities like emotion, motivation, reward or valence processing [59] may like to consider the effect of inter-montage and intra-individual variations in induced EFs inside these nodes and their corresponding network on neural and behavioral outcomes based on the analysis pipeline and the initial findings we have introduced in this study.

## 5. Limitations and future direction

Our study has some limitations that could be addressed in future research. The main limitation is that our results are purely based on simulation. We did not consider other outcome measures such as neural activity/connectivity or behavioral outcomes. We considered averaged EF as an indicator of stimulation intensity. Open questions remain about how to relate EFs with behavioral or neurophysiological changes. The relationship between EF intensity and neural/behavioral responses is not a trivial matter and there is still no clear understanding of the underlying tDCS mechanism of action at the network-level [60]. Integrating EF modeling and neuroimaging information such as functional connectivity that incorporates the interactions within and between large-scale functional networks can provide for a better understanding of tDCS effects in terms of dose-response relationships. For example, Esmaeilpour et al. reported a significant correlation between EF obtained from CHMs and changes in fMRI signals [60]. Kasten et al. revealed a direct link between EF variability and alpha band frequency changes as a proxy indicator of neural activity in a transcranial alternative current stimulation (tACS) study combined with MEG [23]. Combining CHMs with neuroimaging data provide critical insights about tDCS mechanistic effects. However, it is still in an early stage of development and more analysis is needed to increase our understanding of induced EFs and tCDS outcomes in general.

Furthermore, we did not consider network interaction in our analysis pipeline. To our best knowledge, there is no published attempt to find the relationships between induced EFs and interaction between networks that have a critical role in cognitive processes yet. For instance, previous tDCS studies in the treatment of addictive disorders showed potential effects by targeting networks known to be dysregulated in SUDs such as ECN for executive control and DMN for internal ruminations [61]. These two functional networks typically act in counterbalanced order. The hypothesis is that by applying anodal stimulation to DLPFC the level of activity/connectivity should be increased in ECN while it should be decreased in DMN [62]. The results from an fMRI study conducted by Shahbabie et al. demonstrated that application of bilateral DLPFC with the anode electrode over right DLPFC (F4-F3) decreases functional connectivity in DMN especially in posterior parts of the network including middle temporal gyrus, right precuneus, and right PCC [54]. As represented in Fig.7, our calculations showed that temporoparietal and PCC nodes in DMN receive a small amount of EFs in F4-F3 montage. Integrating CHMs with functional data at the network level can shed light on how large-scale functional networks interact with each other by applying tDCS.

Although we calculated EFs in the main nodes of the networks, it is remained unclear which nodes outside the DLPFC are actually important for enhancing tDCS effects on intervention outcomes. The contribution of EF intensity in each part of the network with tDCS outcomes is ambiguous and there is not enough evidence to conclude that which nodes should be stimulated, and which ones should be inhibited to enhance stimulation outcome measures. More physiological and behavioral information is needed to find a relationship between the main components of the networks and tDCS inter-individual responses. This unresolved question would be critical, especially for optimal dose selection and dose customization.

Furthermore, in this study, we only simulated conventional large electrode pades to determine induced EFs at the network-level. Focal stimulation of the network nodes in a group of subjects by using conventional electrodes is difficult. Our study suggests that the probability of interaction between large-scale networks should be considered more carefully using conventional tDCS. Stimulating many nodes of different networks because of the diffusivity of the current makes complexities in modeling the network level contribution in the conventional tDCS montages. Using protocols that produce more focal stimulation such as high-definition (HD) [24], or multi-array electrodes could potentially provide the possibility to control modulatory effects in the network level [37]. In this context, simulation of the HD and multi array electrodes might bring new insights for network level analysis of CHMs.

Another limitation is that our simulations are focused on EF intensity (magnitude of the EF) inside the networks, which is informative of the EF strength. In the current study, we ignored EF components radial to the cortical surface (normal component of the EF), which is informative of EF direction. The normal component of the EF, which is perpendicular to the cortical surface, reflects injected current either entering or leaving the cortex. Previous researches suggest that this component causes polarity-specific effects including facilitatory (anodal like) or inhibitory (cathodal like) effects [5].

In this study, we’ve used a standard functional atlas parcellation to extract large-scale brain networks from personalized CHMs however, atlas-based parcellation provides information from a group of subjects that may be different from the person’s resting-state functional networks. Parcellation of CHMs based on the individual’s functional connectivity might be more accurate to determine EFs inside the main nodes of large-scale-networks. Furthermore, we have only focused on the organization of large-scale distributed networks that obtained from the atlas of resting-state (task-independent) functional networks. Calculation of the EFs according to task-evoked connectivity was not considered in the present study. Investigation of the induced EFs over task-based networks may help to target central brain regions that play pivotal roles in the task performance. For example, a previous study showed that under anodal tDCS (F7-Fp2 montage), during a word generation task, changing connectivity in a given task-related network provides the basis for enhanced neural efficiency in highly specific brain areas critical for task performance. These areas might be located under the stimulation site or distant connected regions [63]. These findings suggest that induced EFs in the major hubs of the task-related network may help understand the mechanistic effects of tDCS. Moreover, targeting main hubs in the task-related networks and functionally connected regions using CHMs may increase the effectiveness of tDCS during task performance since it is hypothesized that tDCS acts on the networks and brain regions concurrently active.

EF modeling in this study is performed based on many assumptions. We used previously established isotropic conductivities for tissues, as is common in the computational studies. Suh et al. reported that the anisotropic skull and WM conductivities significantly affect EF distribution patterns [64]. However, Shahid et al. revealed that anisotropy does not modulate significant changes in comparisons across montages [65]. As mentioned in another study, modeling anisotropy is important when considering deeper target regions inside the brain [66]. Furthermore, our tissue conductivities were completely static, and we did not consider the effects of applying current on tissue conductivities. As mentioned in [42], electrode, tissue, and their interfaces are not merely resistive and can be changed by the applied current intensity that may vary over time. However, these nonlinearity effects have not yet been considered in tDCS modeling studies so far.

Next, we used automatic tissue segmentation and outcomes are strongly dependent on the quality and resolution of T1 and T2-weighted images. However, the quality of segmentation results was evaluated visually slice-by-slice to ensure correct tissue classifications. Furthermore, warping from native space to standard fsaverage space may cause a certain degree of information loss, but it’s necessary for group-level analysis.

Future work is needed to determine whether induced EFs in the main nodes of the large-scale brain networks link to changes in neural activities. It should be investigated that when a single brain region is targeted for stimulation, how other nodes of that region’s network and main components of the other networks interact with the targeted area. Combining CHMs with neuroimaging and behavioral data provides potential insights for understanding how induced EFs interact with the functional activity of the brain at the network level and when stimulation becomes more effective based on current density inside the main components of the networks. Our simulation results from a group of participants with MUDs could be integrated with their clinical and physiological data to determine interindividual variations. Additionally, it would be possible to propose a customized montage by considering the initial state of each individual (e.g. functional connectivity or behavioral measurements at the baseline) and distribution of the current in personalized CHMs. Such effects have not yet incorporated into previous brain stimulation modeling studies.

## 6. Conclusion

In this study we showed that how current flows through the large-scale networks based on group-level analysis of CHMs. In addition to inter-individual variability, we showed that the spread of the EFs in main nodes of the large-scale networks was significantly different between two similar electrode montages for DLPFC stimulation and cathode location can change EF distribution patterns at the network-level especially in the limbic system. The calculation of EFs in the networks and main nodes would be informative since brain stimulation is a network phenomenon. Knowledge about functional connectivity within and between large-scale networks and the EF distribution inside the brain networks might help to resolve ambiguity about how much tDCS effects spread across the brain. The proposed method in this study suggests that a network parcellation of CHMs at the group-level can be used in future studies to understand how tDCS affects large-scale networks and how the results might vary depending on inter-individual variations and electrode arrangements.

## Author Contributions

G.S., H.E., M.S., and F.T. designed the study. H.E., and R.K. collected the data. G.S. performed simulations and data analysis under H.E., M.S., and F.T. supervision. G.S. wrote the paper with input from M.S., M.B., F.T., R.K., M.P.P, and H.E. All authors (G.S., M.S., M.B., F.T., R.K., M.P.P, and H.E.) contributed in manuscript preparation. All authors (G.S., M.S., M.B., F.T., R.K., M.P.P, and H.E.) agreed on the final manuscript before submission.

## Supplementary information

**Table S1:** Group-averaged electric field intensity in [V/m] at large-scale brain networks

| Network | Hemisphere | Average (mean ± SD) EF [V/m] | | P value<br>(Corrected) |
| --- | --- | --- | --- | --- |
|  |  | F4-Fp1 | F4-F3 |  |
| Vis | Left | 0.0523 ± 0.01 | 0.0473 ± 0.01 | **0.0002\*** |
| | Right | 0.0532 ± 0.01 | 0.0488 ± 0.01 | **0.0026\*** |
| SomMot | Left | 0.0972 ± 0.02 | 0.1016 ± 0.02 | 0.1765 |
| | Right | 0.1014 ± 0.02 | 0.1017 ± 0.02 | 0.9284 |
| DorsAttn | Left | 0.0802 ± 0.02 | 0.0873 ± 0.02 | **0.0021\*** |
| | Right | 0.0896 ± 0.02 | 0.0885 ± 0.02 | 0.7708 |
| VentAttn | Left | 0.1257 ± 0.03 | 0.1217 ± 0.03 | 0.3031 |
| | Right | 0.1287 ± 0.03 | 0.1228 ± 0.03 | 0.1883 |
| Limbic | Left | 0.1955 ± 0.04 | 0.1540 ± 0.03 | **0.0000\*** |
| | Right | 0.1552 ± 0.03 | 0.1544 ± 0.03 | 0.8877 |
| ECN | Left | 0.1448 ± 0.03 | 0.1354 ± 0.03 | 0.0550 |
| | Right | 0.1529 ± 0.04 | 0.1476 ± 0.04 | 0.3353 |
| DMN | Left | 0.1500 ± 0.03 | 0.1411 ± 0.03 | 0.0550 |
| | Right | 0.1448 ± 0.03 | 0.1431 ± 0.03 | 0.2786 |

\*Significant differences based on unpaired t-test after FDR correction

**Table S2:** Group-averaged electric field intensity in [V/m] in main nodes of large-scale brain networks

| Network | Node | Hemisphere | Average (mean ± SD) EF [V/m] | | P value<br>(Corrected) |
| --- | --- | --- | --- | --- | --- |
|  |  |  | F4-Fp1 | F4-F3 |  |
| VentAttn | ParOper | Left | 0.0817 ± 0.02 | 0.0785 ± 0.02 | 0.1878 |
| | TempOccPar | Right | 0.0872 ± 0.02 | 0.0807 ± 0.01 | **0.0157\*** |
| | FrOperIns | Left | 0.1514 ± 0.03 | 0.1385 ± 0.03 | **0.0034\*** |
| | | Right | 0.1583 ± 0.03 | 0.1490 ± 0.03 | **0.0479\*** |
| | PFC | Left | 0.2336 ± 0.05 | 0.2154 ± 0.05 | **0.0242\*** |
| | Med | Left | 0.1061 ± 0.1 | 0.1109 ± 0.1 | 0.1593 |
| | | Right | 0.0995 ± 0.02 | 0.0986 ± 0.1 | 0.7921 |
| Limbic | OFC | Left | 0.2474 ± 0.05 | 0.1935 ± 0.04 | **0.0000\*** |
| | | Right | 0.2114 ± 0.04 | 0.2055 ± 0.04 | 0.3533 |
| | TempPole | Left | 0.1342 ± 0.03 | 0.1073 ± 0.02 | **0.0000\*** |
| | | Right | 0.1104 ± 0.02 | 0.1158 ± 0.02 | 0.0631 |
| ECN | TempPar | Left | 0.0635 ± 0.01 | 0.0704 ± 0.02 | **0.0005\*** |
| | | Right | 0.0732 ± 0.02 | 0.0712 ± 0.01 | 0.4020 |
| | PFC | Left | 0.1937 ± 0.04 | 0.1661 ± 0.04 | **0.0000\*** |
| | | Right | 0.1728 ± 0.04 | 0.1650 ± 0.04 | 0.4020 |
| | PcunCing | Left | 0.0688 ± 0.01 | 0.0544 ± 0.01 | **0.0000\*** |
| | | Right | 0.0650 ± 0.01 | 0.0537 ± 0.01 | **0.0000\*** |
| DMN | TempPar | Left | 0.0902 ± 0.02 | 0.0783 ± 0.01 | **0.0000\*** |
| | | Right | 0.0935 ± 0.02 | 0.0884 ± 0.02 | 0.0557 |
| | PFC | Left | 0.2019 ± 0.05 | 0.1966 ± 0.05 | 0.4249 |
| | | Right | 0.2000 ± 0.05 | 0.1985 ± 0.05 | 0.7978 |
| | PcunPCC | Left | 0.0595 ± 0.01 | 0.0500 ± 0.01 | **0.0000\*** |
| | | Right | 0.0629 ± 0.01 | 0.0490 ± 0.01 | **0.0000\*** |

\*Significant differences based on unpaired t-test after FDR correction

## Notes

### Competing Interest Statement

The authors have declared no competing interest.

